# Inhibition of Gasdermin D by Disulfiram Attenuates Cardiac Inflammation and Fibrosis following Ischaemia Reperfusion Injury

**DOI:** 10.64898/2026.03.10.710794

**Authors:** Judy S Choi, Mehnaz Pervin, Helen Kiriazis, Parvin Yavari, Aascha Brown, Man KS Lee, Andrew J Murphy, Daniel Donner, James E Vince, Arpeeta Sharma, Judy B. de Haan

**Author notes:** Co-senior (Arpeeta Sharma, Judy B. de Haan). **Addresses for correspondence: Prof. Judy de Haan,** Lab Head (Cardiovascular Inflammation and Redox Biology Laboratory), Heart Failure Program, Baker Heart and Diabetes Institute, 75 Commercial Road, Melbourne, Victoria 3004, Australia.

## Abstract

**Introduction:** Inadequately controlled inflammation is a key driver of adverse cardiac remodelling after acute myocardial infarction (AMI). Central to this process is activation of the NLRP3 inflammasome-gasdermin D (GSDMD) pathway, which promotes pyroptosis and the release of the pro-inflammatory cytokine interleukin-1β (IL-1β), a mediator strongly associated with infarct severity and poor clinical outcomes. This study investigates whether repurposing the FDA-approved therapeutic Disulfiram, recently shown to inhibit GSDMD pore formation, could reduce inflammation and thus improve cardiac injury after AMI.

**Methods and Results:** Cardiac ischemia-reperfusion (I/R) injury was induced in C57BL/6 mice by 60-minute ligation of the left coronary artery followed by reperfusion. Disulfiram (25 or 50 mg/kg) was administered at reperfusion and daily thereafter. Cardiac function was assessed by echocardiography, while fibrosis and inflammation were evaluated by histology, RT-PCR, immunohistochemistry and immunoblotting. Leukocyte populations in blood, spleen, bone marrow and heart were analysed by flow cytometry. In vitro, mouse bone marrow-derived macrophages (BMDMs) and PMA-differentiated THP-1 cells were treated with Disulfiram. Cytokine secretion, inflammatory gene expression and changes in cell viability (propidium iodide (PI) staining and lactate dehydrogenase (LDH) release) were measured. Disulfiram (50 mg/kg) significantly improved cardiac function 7 days post-I/R. This was accompanied by a significant reduction in cardiac fibrosis and inflammation, as reflected by a lower abundance of inflammatory cells in circulation and cardiac tissue. In LPS- and ATP/Nigericin-stimulated BMDMs and THP-1 cells, Disulfiram dose-dependently (0.1–50 µM) reduced IL-1β and IL-6 secretion and attenuated membrane permeability and cell lysis.

**Conclusions:** This study demonstrates that Disulfiram improves cardiac function post-AMI by ameliorating inflammation and fibrosis, which was associated with reductions in cytokine release from inflammatory cells *in vitro*. Therefore, targeting GSDMD by “repurposing” the FDA-approved drug, Disulfiram, may represent a novel way to provide cardio-protection post-AMI.

**Graphical Abstract:** 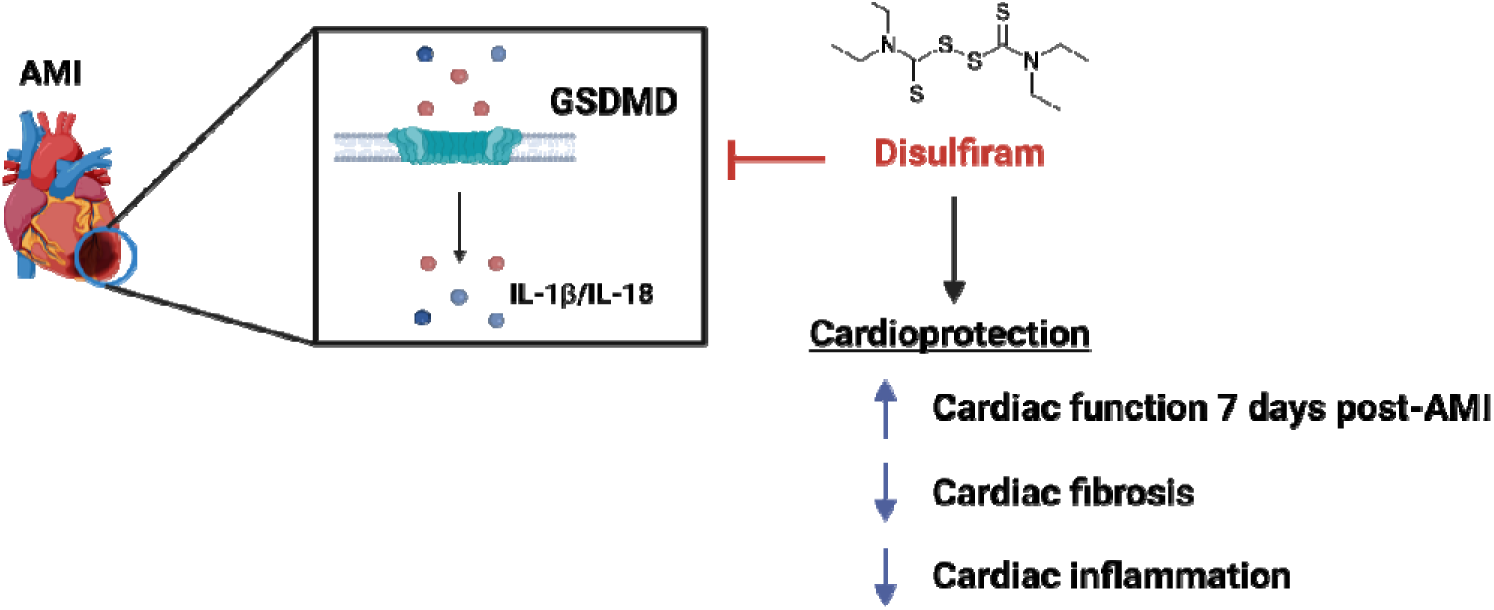

## 1. Introduction

Survival after an acute myocardial infarction (AMI) does not reduce cardiovascular risk, with a substantial proportion of AMI survivors subsequently developing heart failure (HF)^1^. Despite advances in medical techniques to re-oxygenate occluded vessels^2^ and secondary prevention therapeutics that include statins, beta-blockers, angiotensin-converting enzyme (ACE) inhibitors, angiotensin receptor blockers (ARB), mineralocorticoid receptor antagonist (MRA) and sodium glucose cotransporter-2 (SGLT2) inhibitors^3^. Cohort studies indicate that approximately 20-30% of AMI survivors develop HF within the first year which is associated with significantly increased long-term mortality^4^. Post-MI HF represents a high-risk clinical trajectory driven in part by adverse inflammatory and fibrotic remodeling. Furthermore, trends in mortality rates after AMI have undergone minimal change in recent years, highlighting the urgent need for better treatment options to reduce the morbidity and death rate post-AMI^5^. An in-depth understanding of the pathological mechanisms that underpin ischemic injury may lead to novel therapies to lessen the burden of disease and interrupt the progression from AMI to HF.

Inflammation, via activation of the innate immune system, plays a critical role in determining infarct size and cardiac remodelling post-AMI^6, 7^. After an AMI, cellular injury and cardiomyocyte death initiate a rapid pro-inflammatory response. This is characterised by neutrophil infiltration of the myocardium within 12 hours post AMI and a temporal peak at approximately 24 hours as necrotic debris is cleared. This is followed by monocyte-derived macrophage infiltration, peaking at 72 hrs, to phagocytose dead cells^8, 9^. Importantly, the continued presence of inflammatory macrophages in the days to weeks following an AMI drives adverse cardiac remodelling and the progression toward HF^10, 11^. Specifically, persistent leukocytes heighten the inflammatory response by activating the NLRP3 inflammasome and secreting inflammatory cytokines such as IL-1β and IL-18, thereby aggravating scar formation and causing ventricular remodelling^8, 9, 12^.

The NLRP3 inflammasome controls the activation of the cytokines IL-1β and IL-18, and it also induces a highly inflammatory form of cell death, termed pyrotosis, mediated by the formation of GSDMD pores in the cell membrane. Recent evidence suggests that targeting GSDMD to inhibit its pore-forming ability and limit its ability to release damaging cytokines, may offer a novel therapeutic approach to limit post-AMI I/R injury and cardiac remodelling. In support of this notion, Jiang et al. demonstrated that lack of *Gsdmd*, and thereby pore formation, attenuated IL-1β release from CD11b+ cells isolated from the myocardium prevented neutrophil and monocyte infiltration into the infarcted heart, thereby reducing infarct size and improving cardiac function post-AMI^9^. Furthermore, immune cell-specific *Gsdmd* deficiency improved cardiac function and reduced scar size 30 days after permanent ligation-induced AMI^8^. These studies add to the groundswell of support for targeting innate inflammatory responses to improve CVD outcomes.

Disulfiram (DSF), an FDA-approved drug to treat chronic alcoholism, has recently been shown to inhibit GSDMD pore formation through covalent modification of the cysteine residue within the N-terminal region of GSDMD; specifically the Cys191 residue of human and the Cys192 residue of mouse GSDMD^13^. This important discovery highlighted that treatment with cysteine-modifying compounds offers a unique therapeutic approach to lessen pyroptosis-mediated inflammation. A recent study by Zhuang et al., demonstrated that DSF attenuates cardiac injury post-AMI by inhibiting S-palmitoylation of GSDMD at Cys192, thereby impairing membrane localisation of the pore-forming N-terminal fragment and limiting pyroptosis^14^. While these findings revealed an important post-translational regulatory mechanism governing GSDMD activation, the study did not comprehensively interrogate upstream inflammatory signalling pathways. Moreover, the use of a permanent coronary artery ligation model imposes a more severe injury than is typically encountered in patients undergoing percutaneous coronary intervention (PCI) and therefore does not fully replicate ischemia-reperfusion-driven pathology. In the present study, we tested the hypothesis that DSF, by inhibiting GSDMD pore formation and pyroptosis, modulates inflammatory pathways to confer cardioprotection in a clinically relevant model of cardiac ischemia-reperfusion injury. Mechanistically, we focused on the effect of DSF on innate immune responses and its ability to target NLRP3-mediated inflammation to improve cardiac fibrosis. We also hypothesised that inhibiting GSDMD would lead to an improvement in cardiac function in this clinically relevant model.

## 2. Methods

### 2.1 Ethics

All animal experiments were approved by the Alfred Medical Research and Education Precinct animal ethics committee (Ethics number: E8190/2022/B), Melbourne, Australia, and investigations conformed to National Health and Medical Research Council (NHMRC; Australia) guidelines.

### 2.2 *In vivo* ischaemia and reperfusion injury

To study the effect of GSDMD inhibition post-AMI, C57BL/6 male mice were anaesthetised with isoflurane and baseline echocardiography was performed. At 12-weeks of age, animals were randomly allocated to either the sham or Ischaemia/reperfusion (I/R) group. Animals were anaesthetised with an intraperitoneal (IP) injection of ketamine, xylazine and atropine mixture (KXA; 100mg/kg of ketamine, 20mg/kg of Xylazine and 1.2mg/kg of Atropine). Animals were intubated with a tracheal cannula connected to a ventilator to assist with breathing during surgery. The left coronary artery (LAD) was ligated using a 7-0 silk suture and the chest cavity and skin layers were subsequently closed with 6-0 monofilament sutures. Animals allocated to the sham group did not have the left coronary artery ligated, and the chest cavity and skin layers were closed immediately after the heart was exposed. Animals underwent 1 hour of ischemia and were administered DSF (25 or 50mg/kg) or a vehicle (sesame oil) via IP injection immediately at reperfusion. DSF treatment was administered daily thereafter until termination. Echocardiography was performed either at 7 days (early cardiac remodelling) or 28 days post-AMI (late cardiac remodelling) to assess changes in cardiac function.

### 2.3 Echocardiography

Echocardiograms were obtained on a Vevo F2 ultrasound scan system with UHF57x transducers. Mice were anaesthetised with 4.8% isoflurane gas in an induction box and transferred to a nose cone with 1.8% isoflurane for the duration of the procedure. Echocardiograms were analysed using Vevo LAB software (Fujifilm VisualSonics). An internal quality control process was performed on all measurements in a blinded manner. Two-dimensional B-mode long-axis left ventricle echocardiogram images were used to evaluate changes in systolic function. The left ventricle endocardium was traced at the end-diastole and end-systole phases to calculate ejection fraction, longitudinal fractional shortening, cardiac output, stroke volume, systolic and diastolic volume and area. Two images were analysed and averaged per mouse.

### 2.4 Tissue Collection

At termination, mice were euthanised by CO_2_ asphyxiation and blood was drawn (∼1mL) into a heparinised syringe and plasma separated for cytokine analysis. The left ventricle was dissected into three parts: base, mid-ventricular ring and apex. The base of the heart was snap-frozen for gene expression analysis, and the mid-ventricular ring was fixed in 10% neutral buffered formation (NBF) and paraffin-embedded for histological analysis. The apex of the hearts from the 7-day post-AMI studies were used for flow cytometry to analyse changes in immune cell populations, and apical tissues from the 28-day post-AMI studies were snap-frozen in liquid nitrogen for protein expression analysis. The spleen, blood and bone marrow were processed to analyse changes in immune cell populations using flow cytometry.

### 2.5 Histology

Paraffin embedding, sectioning, mounting and basic histological staining (Picrosirius Red and Masson’s trichrome) of the left mid-ventricular ring sections were performed by the Monash Histology Platform. For picrosirius red staining, images were captured by an Olympus BX43 light microscope (Olympus) in brightfield and polarised light at 20x magnification. Positive staining was quantitated with Image-Pro Analyzer 7.0 (Media Cybernetics).

Paraffin-embedded heart tissues were stained for CD68, a marker highly expressed by monocytes and circulating and tissue-resident macrophages, to evaluate macrophage infiltration into the heart as described previously^15^.

### 2.6 Flow cytometry

The immune cell populations in the heart, bone marrow, spleen and blood were examined by staining with immune cell markers of neutrophils, monocytes and macrophages and analysing the cell populations by flow cytometry as described previously^16^. Samples were analysed using FlowJo (v10.10; BD Biosciences).

### 2.7 Immunoblot analysis

Protein extraction was performed by homogenizing left ventricular apical tissue in 10µL ice-cold DISC lysis buffer containing complete protease inhibitor and Phos-STOP phosphatase inhibitor per mg of tissue, using a Tissue Lyser 85300. Protein concentration of the lysates was quantified with a Pierce™ BCA protein assay kit (Thermo Fisher, #23227). Proteins were separated by electrophoresis in NuPage 4-12% Bis-Tris gels and transferred to PVDF membranes. Membranes were probed with goat anti-IL-1β (R&D, #AF-401-NA), mouse anti-NLRP3 (Adiopogen, #AG-20B-0014-C100), rabbit anti-GSDMD (Abcam, #AB209845), rabbit anti-caspase-1 (Abcam, #AB179515), rabbit anti-caspase-3 (Cell Signalling, #9662) and rabbit anti-GAPDH (Cell Signalling, #5174) antibody for equal loading verification. Signals were enhanced with Immobilon^®^ Forte Western HRP Substrate (Millipore, #WBLUF0500) chemiluminescent and the membranes were then scanned using Bio-rad G-Box (Biorad).

### 2.8 Gene expression analysis

Total RNA was extracted using TRIzol after homogenisation of snap frozen left ventricular base tissue. Real-time quantification of gene expression (Supplementary Table 4) was performed and was analysed as described previously^15^.

### 2.9 Cell culture

Primary bone marrow–derived macrophages (BMDMs), isolated from C57/BL6 mice, were grown in l-cell conditioned media and THP-1 monocytes were cultured as per our previous study^15^. Briefly, BMDMs and THP-1 monocytes were primed with lipopolysaccharide (LPS) (4h; 0.1 μg/mL) and treated with DSF (3h; 0.1-50µM) and activated with ATP (1h; 1 mmol/L). For gene expression studies, cells were primed with LPS (3h; 0.1 μg/mL), treated with DSF (3h; 0.1-50µM). IL-1β and IL-6 secretion in the supernatant was determined by ELISA as per the manufacturer’s instructions. RNA extraction, cDNA synthesis, and qRT-PCR were performed as described previously^15^.

### 2.10 Statistical analysis

All statistical analyses were conducted using GraphPad Prism (Version 10.6.1). Outlier testing was conducted using the ROUT test followed by Shapiro-Wilk normality tests to confirm their normal distribution. Results that did not meet the normality assumption were subsequently analysed using a non-parametric Kruskal-Wallis test for significance. Results that were normally distributed with equal variance were analysed using one-way ANOVA followed by a Tukey’s multiple comparisons test. For results that were normally distributed but had non-equal variance, the Brown-Forsythe and Welch one-way ANOVA test was applied, followed by Dunnett’s multiple comparison test. All results are expressed as mean ± SEM and are considered statistically significant if P < 0.05.

## 3. Results

### 3.1 Disulfiram attenuates inflammation in BMDMs

Cultured WT BMDMs that are primed with LPS (Figure 1A and 1C) and NLRP3 activated with ATP (Figure 1B and 1D) demonstrate significantly increased *Il-1β* and *Il-6* gene expression and protein secretion respectively. *Il-1β* and *Il-6* gene expression remained unchanged in LPS-treated *Gsdmd^-/-^*BMDMs compared to LPS-treated WT BMDMs, however protein secretion of IL-1β and IL-6 was markedly suppressed in LPS+ATP treated *Gsdmd^-/-^* BMDMs compared to the LPS+ATP-treated WT BMDMs (Figure 1B and 1D). Treatment with DSF significantly attenuated IL-1β secretion by up to 95% in a dose-dependent manner in WT BMDMs (Figure 1B), whilst having no effect in *Gsdmd^-/-^* BMDMs, compared to their respective LPS+ATP-treated controls. DSF treatment reduced *Il-1β* gene expression in WT BMDMs and in *Gsdmd^-/-^* BMDMs (Figure 1A) compared to LPS alone. IL-6 protein secretion was reduced by up to ∼91% in a dose-dependent manner in WT BMDMs and similarly by up to ∼90% in *Gsdmd^-/-^*BMDMs (Figure 1D). This effect was also seen at the transcriptional level where DSF treatment reduced *Il-6* gene expression in WT BMDMs and in *Gsdmd^-/-^* BMDMs (Figure 1C) compared to their respective LPS-treated controls. These data confirm that the absence of GSDMD protein and hence absence of pore formation, reduces IL-1β secretion independent of its gene expression. Furthermore, the downstream cytokine, IL-6, was attenuated both at the gene and protein level by DSF, indicating transcriptional suppression of inflammation downstream of IL-1β by DSF.

**Figure 1:**
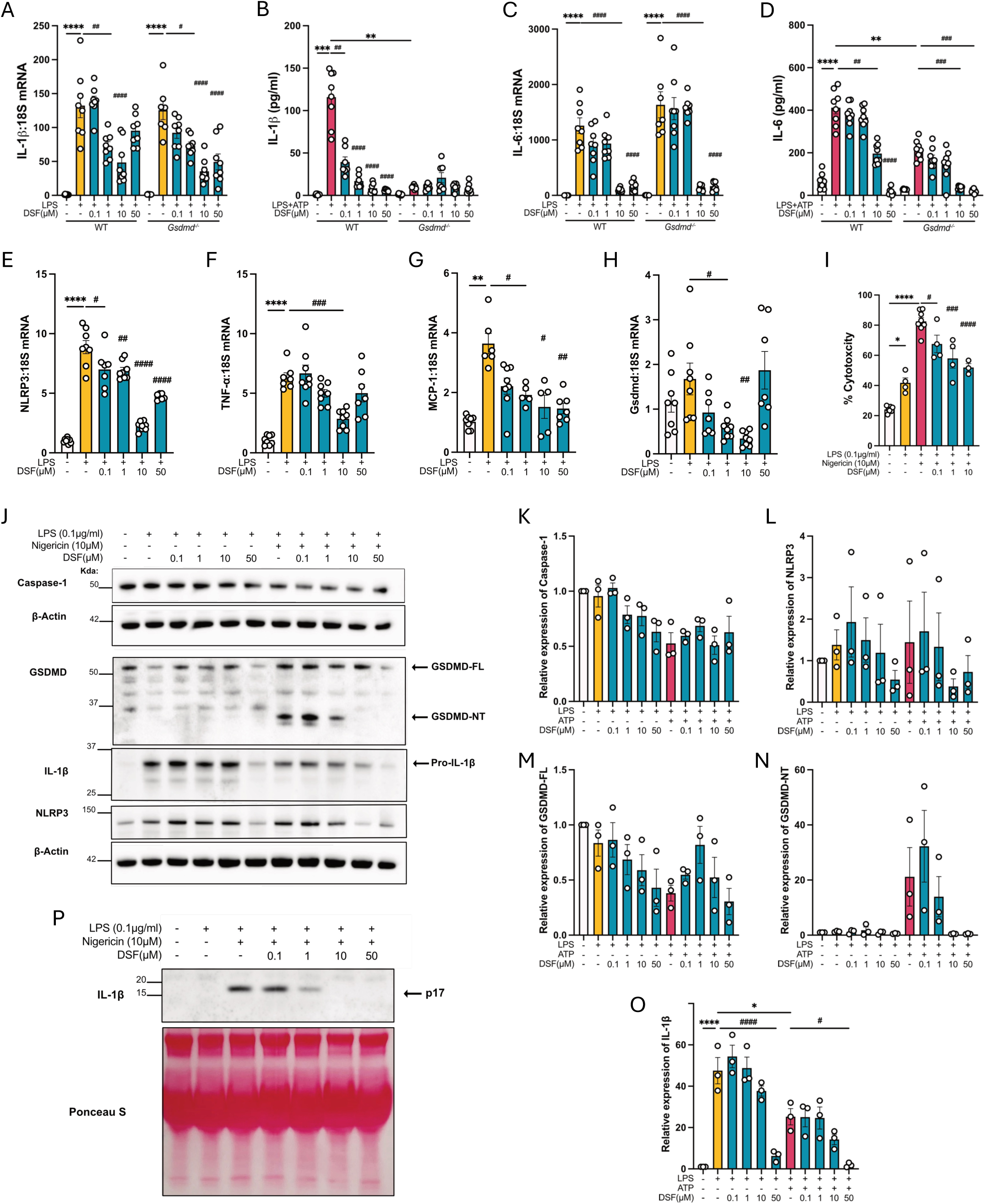
BMDMs were treated with LPS (0.1μg/μL; 4-hours) ± ATP (1mM; 1-hour/3-hour after LPS treatment respectively) and ± DSF (0.01-50μM; 1-hour after LPS treatment). (A) *Il-1β* gene expression, (B) IL-1β secretion, (C) *Il-6* gene expression and (D) IL-6 secretion in WT and *Gsdmd*^-/-^BMDMs. Gene expression of (E) *Nlrp3*, (F) *Tnf-α*, (G) *Mcp-1* and (H) *Gsdmd*. (I) LDH levels as measured in the supernatants to determine cell lysis. (J) Representative immunoblot and analysis of (K) Caspase-1, (L) NLRP3, (M) GSDMD, (N) GSDMD-NT and (O) pro-IL-1β. (P) Representative immunoblot of cleaved IL-1β (p17) in supernatants collected from WT BMDMs. Statistics used: one-way ANOVA. *P<0.05, **P<0.01, ****P<0.0001, ^#^P<0.05, ^##^P<0.01, ^###^P<0.001 and ^####^P<0.0001 as indicated. Results are expressed as mean ± SEM. Data are representative of three biological replicates.

In addition, the effect of DSF was examined on inflammatory gene expression in WT BMDMs. LPS treatment led to a substantial increase in *Nlrp3*, *Tnf-α*, and *Mcp-1* gene expression by 6.4-, 3.6- and 8.7- fold respectively compared to their respective controls (Figure 1E-1G), while DSF treatment attenuated *Nlrp3, Tnf-α, Mcp-1*, and *Gsdmd* gene expression (Figure 1E-1H) compared to LPS treatment.

Loss of membrane integrity results in the release of cytosolic LDH and is indicative of cell viability^17^. Treatment with LPS and LPS + Nigericin led to a 1.7- and 3.4-fold increase in LDH release respectively compared to the untreated control (Figure 1I). LDH release was significantly attenuated by DSF treatment in a dose-dependent manner (Figure 1I).

Protein expression of Caspase-1, GSDMD, IL-1β and NLRP3 was determined by immunoblotting to investigate if DSF treatment influenced key proteins of the NLRP3-inflammasome-GSDMD pathway. Cytosolic levels of Caspase-1 remained virtually unaltered across all treatment groups, suggesting that DSF has no effect on Caspase-1 levels (Figure 1J and 1K). Consistent with the existing literature, both priming with LPS and activation with either ATP or nigericin are required for the cleavage of GSDMD into GSDMD-NT, which is inserted into the plasma membrane to form pores^18^. Treatment with LPS and nigericin resulted in the appearance of the GSDMD-NT (p30) which was absent in LPS-only treated group. GSDMD-FL was mostly unaffected by DSF treatment, only showing a tendency towards a reduction at 50 µM DSF (Figure 1J and 1M). There was a reduction in GSDMD-NT with DSF treatment at 1.0 µM, with virtually undetectable levels at 10 and 50µM (Figure 1J and 1N). Importantly, DSF treatment significantly reduced pro-IL-1β at 50µM when treated with LPS with a tendency toward reduction at 10uM (Figure 1J and 1O). Interestingly, in the presence of LPS and nigericin, the level of pro-IL-1β was significantly lower than that seen in the presence of LPS only (Figure 1J and 1O), with DSF treatment lessening the level of pro-IL-1β after LPS and nigericin treatment at 50µM under both conditions (Figure 1J and 1P). Most importantly, secretion of cleaved IL-1β (p17) was dose-dependently reduced in WT BMDMs treated with LPS and nigericin after DSF treatment (Figure 1P).

Overall, these results suggest that GSDMD pore formation is required for rapid IL-1β secretion from BMDMs, and that treatment with DSF, a known GSDMD pore formation inhibitor, reduces pro-inflammatory gene and protein expression.

### 3.2 Inhibition of GSDMD with disulfiram attenuates inflammation in PMA-differentiated THP-1 macrophages

The effect of DSF treatment was assessed in human macrophages (PMA-differentiated THP-1 cells) providing insight into the inflammatory response of cells involved in human cardiac injury post-AMI and exposed to DSF treatment.

PMA-differentiated THP-1 cells were primed with LPS and stimulated with ATP, which led to a significant increase in IL-1β secretion compared with controls (Figure 2A). The increase in IL-1β secretion was attenuated after DSF treatment at 10µM and 50µM (Figure 2A). LPS treatment alone led to a significant increase in *IL-1β, IL-6, MCP-1* and *TNF-α* gene expression, which was significantly attenuated with DSF treatment (Figure 2B-E).

**Figure 2:**
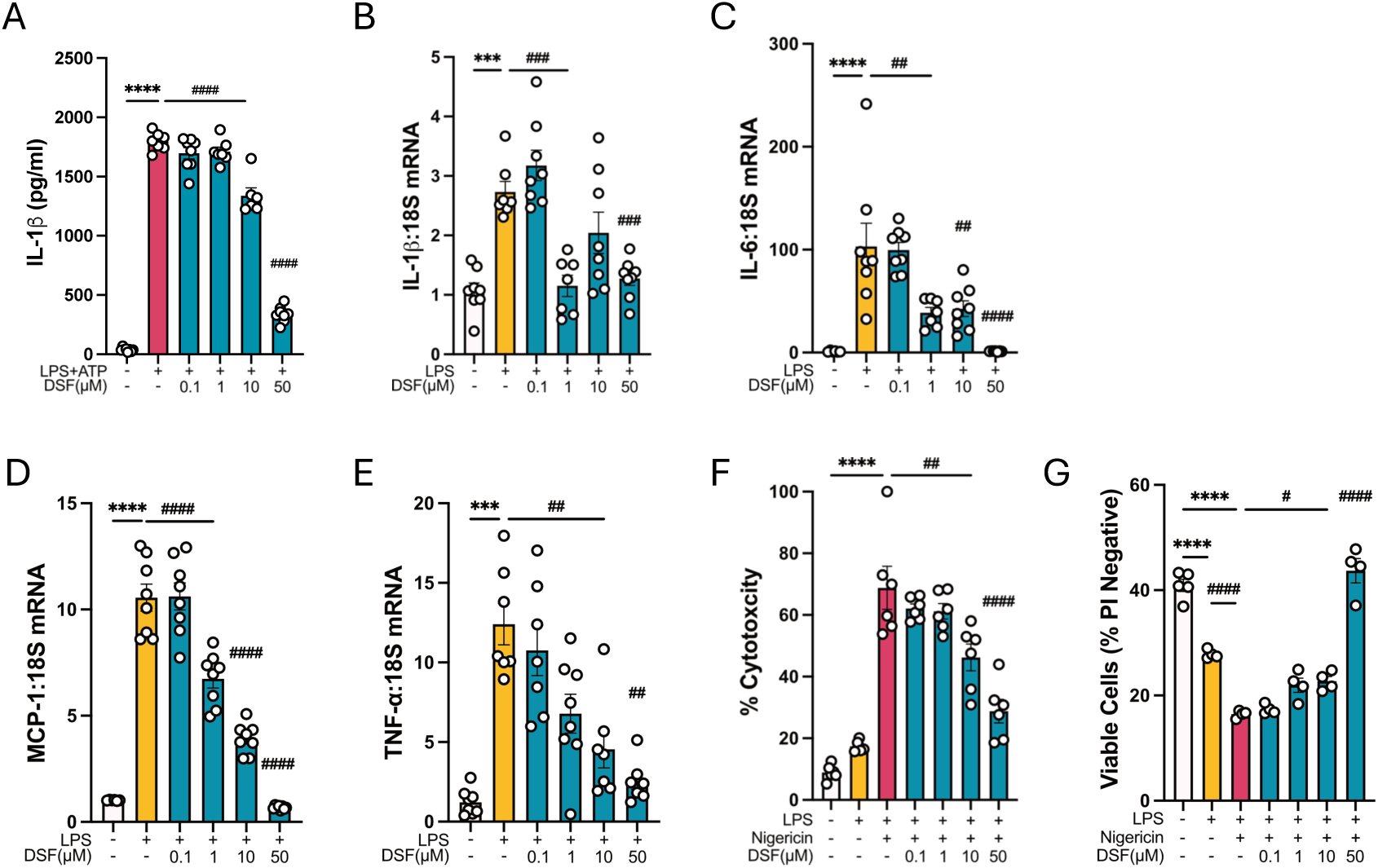
THP-1 cells were treated with LPS (0.1μg/μL; 4-hours) ± Nigericin/ATP (10µM Nigericin/ 1mM ATP; 1-hour/3-hour after LPS treatment respectively) and ± DSF (0.01-50μM; 1-hour after LPS treatment). (A) IL-1β secretion, (B) *IL-1β* gene expression and (C) *IL-6* gene expression in treated THP-1 cells. Gene expression of (D) *MCP-1* and (E) *TNF-α* in treated THP-1 cells. (F) LDH levels and (G) Cell viability (% PI negative) measured from the cell supernatants. Statistics used: one-way ANOVA. ***P<0.001 and ****P<0.0001 as indicated. ^##^P<0.01, ^###^P<0.001 and ^####^P<0.0001 vs LPS+Nigericin unless otherwise indicated. Results are expressed as mean ± SEM. Data are representative of three independent biological replicates.

The effect of DSF on cell viability was assessed by two independent methods. First, LDH release was measured from dying cells and second, cells were stained with the membrane impermeable dye propidium iodide (PI). LDH release was increased by 7.8-fold after LPS and nigericin treatment compared with untreated controls. This increase was significantly attenuated by up to ∼60% with DSF in a dose-dependent manner (Figure 2F). In the cell viability assay, treatment with LPS alone led to a ∼32% reduction in cell viability whilst LPS+nigericin treatment led to a ∼60% reduction in viable cells (Figure 2G) compared to controls. DSF treatment significantly increased cell viability in a dose-dependent manner (Figure 2G).

Overall, these results suggest that DSF attenuates IL-1β secretion as well as the gene expression of a range of pro-inflammatory mediators in human THP-1 cells. Additionally, DSF protects against inflammatory cell death of human macrophages, thereby limiting the release of cytotoxic cytokines.

### 3.3 Disulfiram improves cardiac function and reduces cardiac fibrosis

Mice underwent temporary LAD ligation for an hour to assess the therapeutic value of GSDMD inhibition by DSF (25,50mg/kg) on cardiac remodelling 7 and 28 days post-I/R injury.

Morphometric analysis at endpoint is shown in Supplementary Table 1 (7-day) and Table 2 (28-day). LAD ligation caused no difference in body weight, heart weight and LV weight at both 7 days (Sup Table 1) and 28 days (Supplementary Table 2) post-AMI. A trend towards an increase in the ratio of heart weight/tibia length (Supplementary Table 1) was observed at 7 days post-I/R injury compared to the sham group. This was attended along with respect to the LV weight/tibia length ratio with DSF (50mg/kg) prevented this increase in heart weight/tibia length ratio (Supplementary Table 1). These changes were not observed in the 28 days post-I/R injury cohort (Supplementary Table 2).

At 7-days post-I/R injury, significant changes in almost all parameters were noted in mice that underwent I/R surgery alone relative to sham mice (Figure 3C-H). The I/R group displayed the following characteristics: diastolic volume increased by 1.6-fold reflecting ventricular remodelling and dilation (Figure 3D), systolic volume increased by 2.3-fold (Figure 3E), ejection fraction decreased by 50% (Figure 3G) and longitudinal fractional shortening reduced by 48% (Figure 3H). No differences in heart rate (Figure 3C) and cardiac output (Figure 3F) were observed. DSF (50mg/kg) treatment significantly improved diastolic volume (Figure 3D), systolic volume (Figure 3E) and ejection fraction (Figure 3H) compared to the vehicle-treated I/R group. These data demonstrate that there is ventricular remodelling with inadequate contraction and filling of the heart post-I/R, which was improved by DSF treatment.

**Figure 3:**
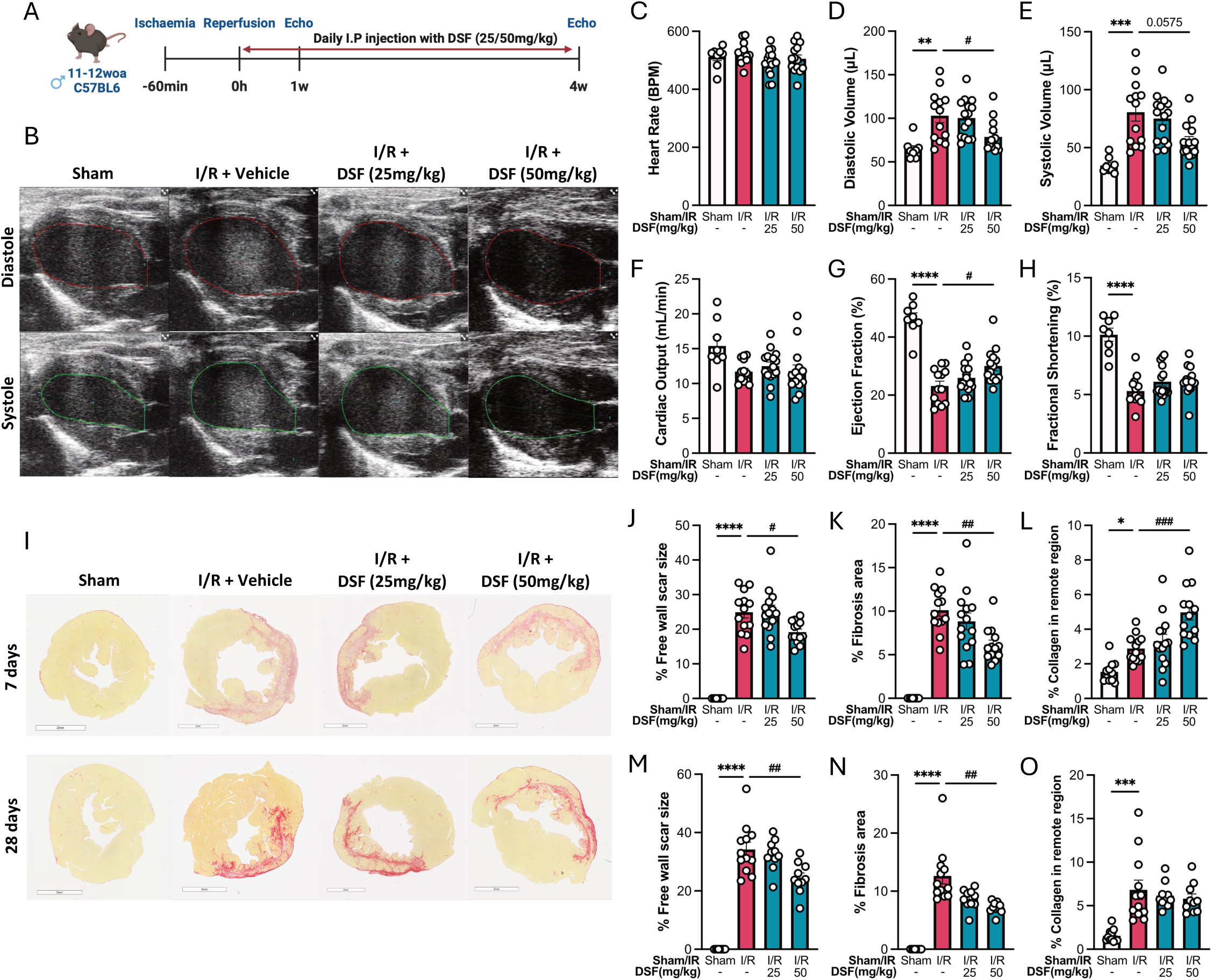
(A) Experimental timeline for I/R study. (B) Representative echocardiographic images at end-phase diastole and end-phase systole. The red lines show the tracing of the endocardium at maximum relaxation (diastole) and the green lines reflect maximum contraction at systole. Analysis of (C) Heart rate, (D) Diastolic volume, (E) Systolic volume, (F) Cardiac output, (G) Ejection fraction and (H) Longitudinal fractional shortening is shown in sham and I/R mice hearts treated with vehicle or DSF 7 days post-I/R injury. (I) Picrosirius red (PSR) stained hearts and quantification of (J) % free wall scar size, (K) % fibrosis area and (L) % collagen in the remote region 7 days post-I/R injury and (M) % free wall scar size, (N) % fibrosis area and (O) % collagen in the remote region 28 days post-I/R injury in the hearts treated with vehicle or DSF. Scale bars represent 2mm. *n* represents the number of biological replicates, which are graphed as individual points. *n*= 8-14. Statistics used: Ordinary one-way ANOVA (D, G, J, M), Brown-Forsythe and Welch ANOVA (E) and Kruskal-Wallis (F, H, K, L, N, O) were performed. **P<0.01, ***P<0.001, ****P<0.0001, ^#^P<0.05, ^##^P<0.01 and ^###^P<0.001. Results are expressed as mean ± SEM.

Cardiac fibrosis (Figure 3I) was measured by evaluating the percentage free wall scar size (the area of fibrosis within the infarct zone) (Figure 3J), the % fibrosis area (Figure 3K) and the percentage of collagen in the remote infarct-free region at 7-days post-AMI (Figure 3L). No fibrosis was observed in the sham group (Figure 3J) 7-days after sham surgery. Vehicle-treated I/R-treated hearts exhibited an increase in free wall scar size (Figure 3J), total fibrosis (Figure 3K), and an increase in collagen in the remote infarct-free region (Figure 3L) compared to their respective sham groups. Importantly, treatment with DSF at 50mg/kg significantly attenuated the free wall scar size (Figure 3J) and overall fibrosis area (Figure 3J) compared to the vehicle-treated I/R group. Interestingly, the percentage collagen in the remote region was elevated 1.7-fold after DSF treatment (50mg/kg) compared to the I/R group at the 7-day timepoint (Figure 3K).

At the later stage of cardiac remodelling (28 days post-I/R injury, Figure 3I), similar to that seen at 7 days post-I/R injury, the % free wall scar size (Figure 3M), the % fibrosis area (Figure 3N) and the % collagen in the remote region (Figure 3O) were all highly significantly increased. DSF treatment (50mg/kg) significantly attenuated both the % free wall scar size and fibrosis area (Figure 3M-3N) but had no effect on the % collagen in the remote region of the heart (Figure 3O). After 28 days, cardiac functional data (diastolic volume (SFigure 1C), systolic volume (SFigure 1D), cardiac output (SFigure 1E), ejection fraction (SFigure 1F), fractional shortening (SFigure 1G) and global longitudinal strain (SFigure 1H) were significantly impaired post-AMI; however, DSF treatment did not improve these parameters (SFigure 1B-H) in contrast to that observed 7 days post-I/R injury (Figure 3D,3E and 3G).

### 3.4 Disulfiram treatment inhibits post-AMI inflammatory response

Cell populations of the bone marrow, spleen, blood (Figure 4) and hearts (Figure 5F-H) were analysed by flow cytometry 7 days post-I/R injury, to evaluate whether DSF treatment alters immune cell populations known to express GSDMD.

**Figure 4:**
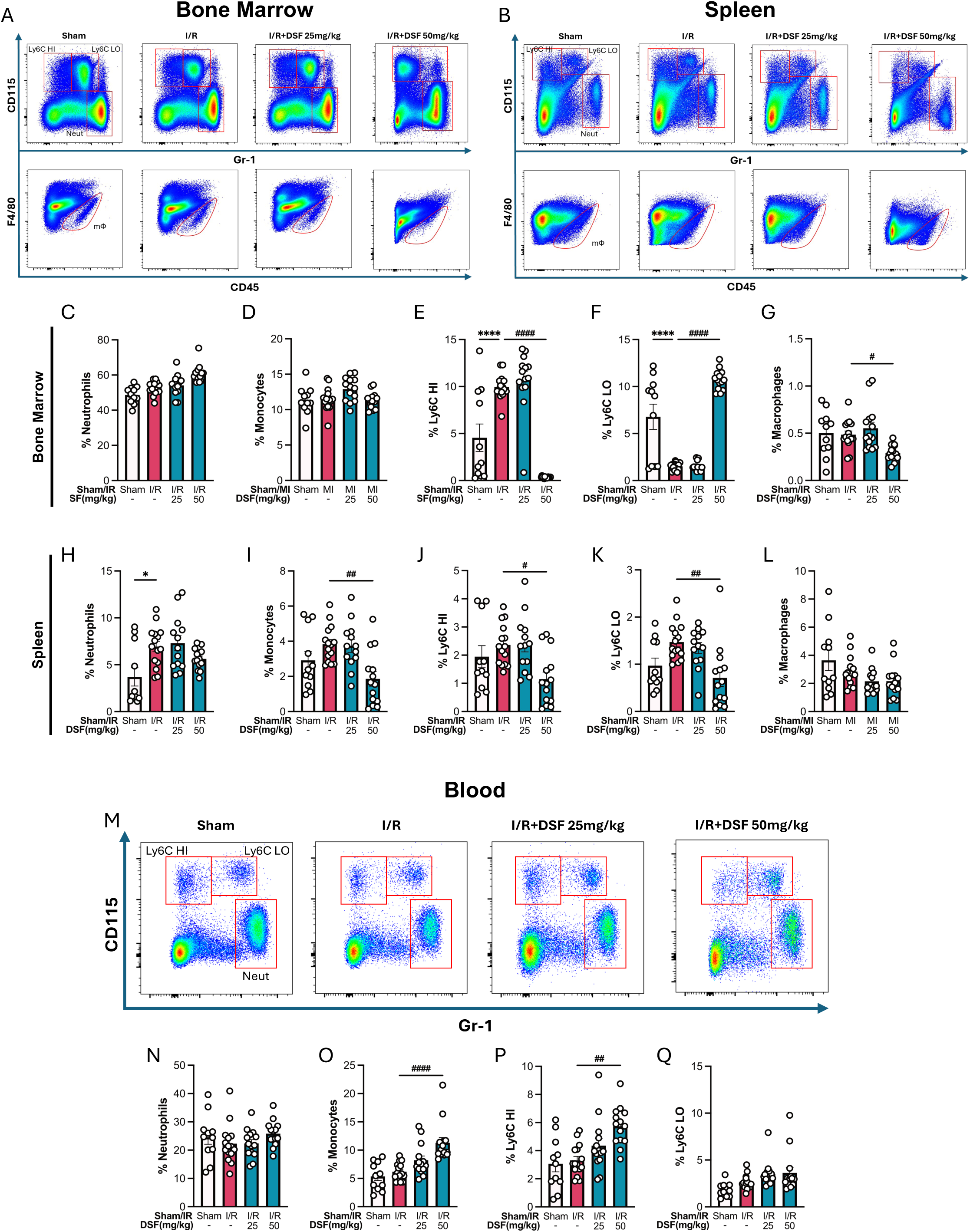
(A) Flow cytometry gating of the (A) bone marrow and (B) spleen from sham and I/R mice treated with vehicle or DSF at 7 days post-I/R injury. Quantification of (C) % Neutrophils, % (D) Total monocytes, (E) % Ly6C-high monocytes, (F) % Ly6C-low monocytes and (G) % macrophages in bone marrow. Quantification of (H) % Neutrophils, (I) % Total monocytes, % (J) Ly6C-high monocytes, %(K) Ly6C-low monocytes and (L) % Macrophages measured in spleen. (M) Flow cytometry gating of the blood from sham and I/R mice treated with vehicle or DSF at 7 days post I/R injury. Quantification of (N) % Neutrophils, (O) % Total monocytes, (P) % Ly6C-high monocytes and (Q) % Ly6C-low monocytes in blood. *n* represents the number of biological replicates, which are graphed as individual points. Statistics used: Brown-Forsythe and Welch ANOVA (D, E, H, I, J, K, P) and Ordinary one-way ANOVA (F). *n*=11-15. *P<0.05, ****P<0.0001, ^#^P<0.05, ^##^P<0.01 and ^###^P<0.001. Results are expressed as mean ± SEM.

**Figure 5:**
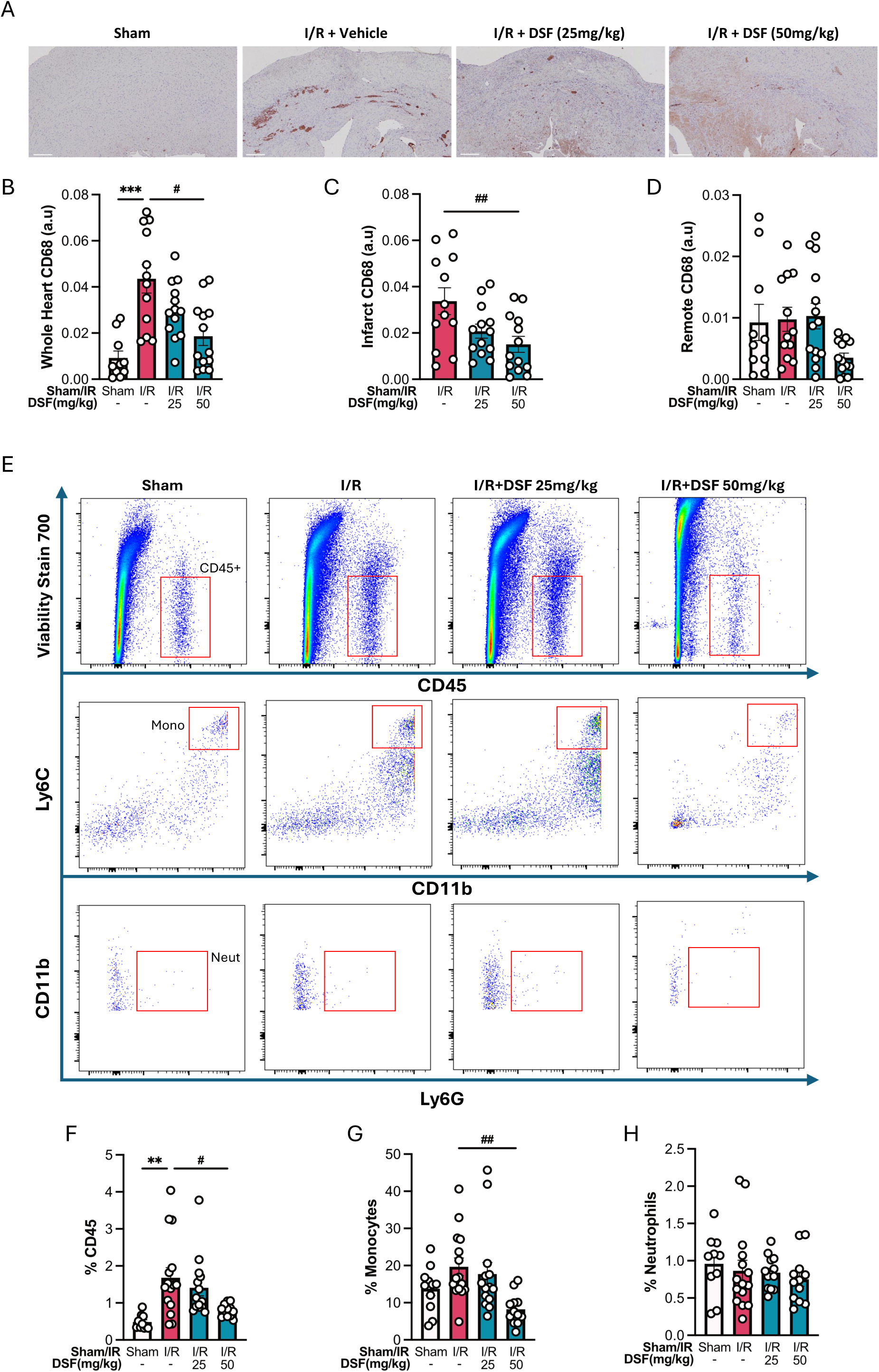
(A) Representative images of CD68 staining in the heart and quantification in the (B) whole heart, (C) infarct region and (D) remote region in sham and I/R mice treated with vehicle or DSF at 7 days post-I/R injury. Scale bars represent 200µm. (E) Flow cytometry gating of the heart from sham and I/R mice treated with vehicle or DSF at 7 days post-I/R injury. Quantification of (F) % CD45^+^ leucocytes, (G) % Monocyte and (H) % Neutrophils in heart. *n* represents the number of biological replicates, which are graphed as individual points. Statistics used: Kruskal-Wallis (B, F, G) and Ordinary one-way ANOVA (C). *n*=10-15. **P<0.01, ***P<0.001, ****P<0.0001, ^#^P<0.05 and ^##^P<0.01. Results are expressed as mean ± SEM.

In the bone marrow, the most significant changes occurred in the monocyte sub-populations. I/R injury caused an increase in pro-inflammatory Ly6C-high expressing monocytes (Figure 4D), while Ly6C-low expressing monocytes were reduced (Figure 4E). Importantly, DSF (50mg/kg) treatment significantly reversed this, resulting in a 96% reduction in Ly6C-high monocytes (Figure 4D) and a 7-fold increase in Ly6C-low monocytes (Figure 4E). Furthermore, macrophages were reduced by ∼40% after DSF (50 mg/kg) treatment (Figure 4F). At this time point, there were no significant differences in the % neutrophils across all groups (Figure 4B). Overall, suggesting that DSF at the higher dose of 50mg/kg caused a phenotypic shift in the monocyte population, resulting in an abundance of anti-inflammatory monocytes, accompanied by reductions in macrophages in the bone marrow.

In the spleen, neutrophils were significantly increased by ∼1.9-fold (Figure 4H) after I/R injury but DSF treatment had no effect on this cell population. DSF (50mg/kg) treatment led to a ∼51% reduction in monocytes (Figure 4I) compared to the untreated I/R group. Sub-population analysis showed a ∼50% reduction in both Ly6C-high and Ly6C-low monocytes after DSF treatment (Figure 4J-4K), while the macrophage population was unaffected by both I/R and DSF treatment (Fig 4L).

In the blood, no change in the percentage of neutrophils and monocytes was observed across all groups (Figure 4N, 4O), although separation into subsets revealed a significant 1.7-fold increase (Figure 4P) in inflammatory Ly6C-high monocytes after 50mg/kg DSF treatment.

### 3.5 Disulfiram attenuates monocyte and macrophage infiltration into the heart post-AMI

Post-AMI, monocytes and macrophages are recruited to sites of injury and robustly secrete the pro-inflammatory cytokine IL-1β, particularly through the activation of the NLRP3 inflammasome-GSDMD axis ^19^ ^20^. In line with current literature, our data shows significantly increased CD68+ throughout the heart tissue 7 days post-AMI (Figure 5B). DSF caused a dose-dependent decrease in CD68+ staining, with a significant ∼57% decrease reached after 50mg/kg DSF treatment in the heart tissue (Figure 5B) with a ∼55% decrease within the infarct region (Figure 5C). At 28 days post-I/R, CD68+ levels were significantly elevated throughout the whole heart section and unchanged by DSF (SFigure 1J), whilst no significant changes were noted after both doses of DSF treatment within the infarct or remote region of the heart (SFigure 1K-L).

Consistent with published literature, flow cytometry of heart tissue revealed an increased number of leukocytes in I/R-injured heart, as shown by a 3.5-fold increase in CD45^+^ cells (Figure 5F) compared to the sham group. No significant differences in neutrophils (Figure 5H) were observed across all groups. Importantly, DSF (50mg/kg) caused a reduction in total leukocyte numbers (Fig 5F) and specifically in the % cardiac monocytes (Figure 5G) compared to the vehicle-treated I/R group.

### 3.6 Effect of disulfiram on cardiac gene expression of inflammatory and oxidative stress markers

Cardiac gene expression was analysed at 7 and 28 days post-I/R injury to determine if the reduction in cardiac fibrosis and improvement in cardiac function observed after DSF treatment are influenced by genes that regulate inflammation, fibrosis and cell death pathways.

At 7 days post-AMI, the inflammatory genes, *Il-6* and *Il-18* were significantly elevated by 6.4- and 2.7-fold (Figure 6A). DSF (50mg/kg) treatment attenuated pro-inflammatory cytokine gene expression of *Il-1β, Il-6, Il-18, Ifn-*γ, and *Tnf-*α (Figure 6A and SFigure 2A). The gene expression of the chemokine *Mcp-1* remained unaltered across all groups. Intriguingly, at 7 days post-AMI, the anti-inflammatory marker, *Il-10*, was significantly attenuated after DSF (50mg/kg) treatment (Figure 6A).

**Figure 6:**
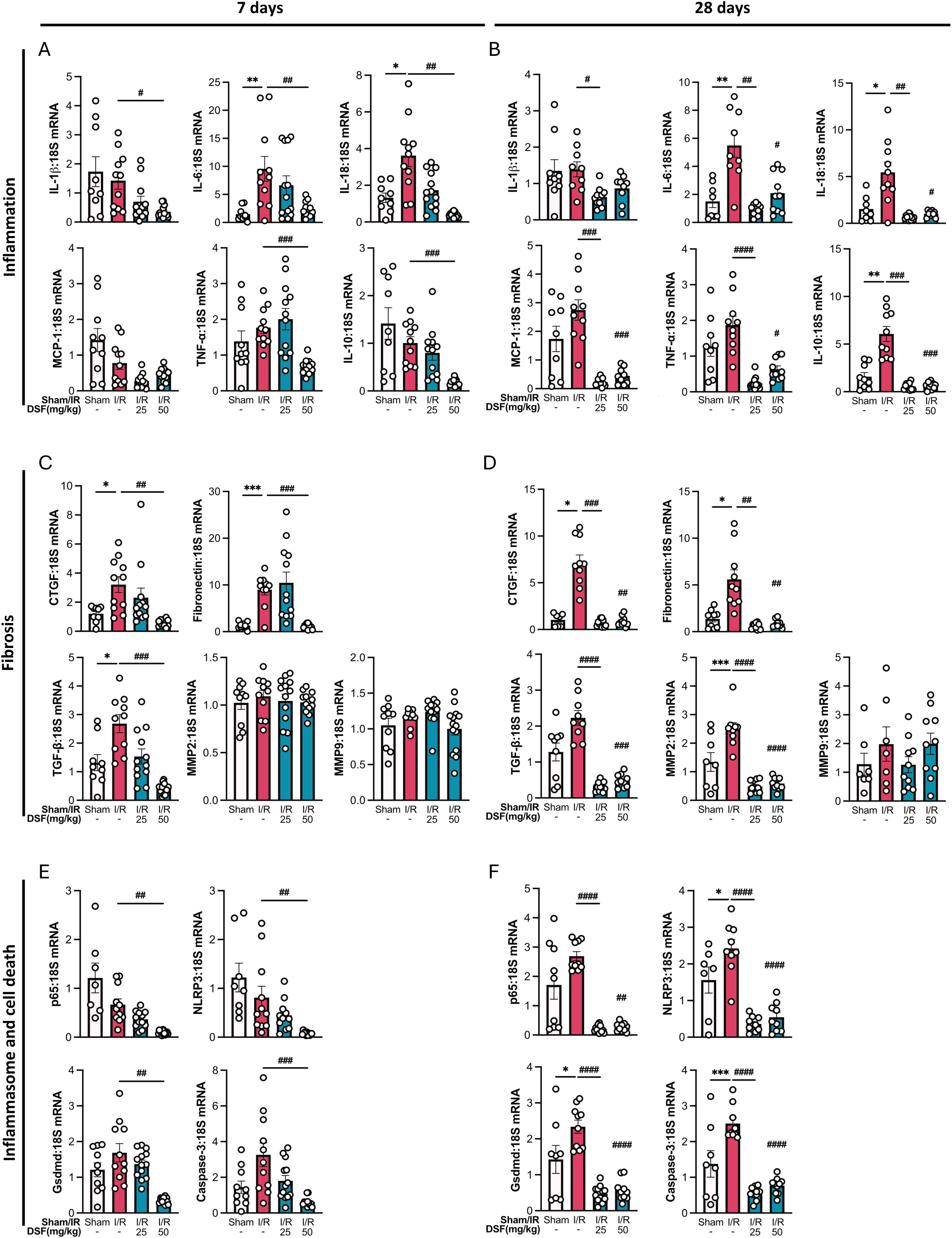
Inflammatory gene expression of *Il-1β, Il-6, Il-18, Mcp-1, Tnf-α and Il-10* in the heart determined by qRT-PCR (A) 7 days or (B) 28 days post-I/R injury with vehicle or DSF. Expression levels of fibrosis genes *Ctgf, Fibronectin, Tgf-β, Mmp2 and Mmp9* in the heart as determined by qRT-PCR (C) 7 days or (D) 28 days post-I/R injury with vehicle or DSF. Inflammasome and cell death-related gene expression analysis of *p65* subunit of NF-κB, *Nlrp3*, *Gsdmd* and *Caspase-3* in the heart determined by qRT-PCR (E) 7 days or (F) 28 days post-I/R injury with vehicle or DSF. Gene expression is shown relative to 18S mRNA levels. *n* represents the number of biological replicates, which are graphed as individual points. Statistics used: Kruskal-Wallis (*Il-1β, Caspase-3* and *Nlrp3*), Ordinary one-way ANOVA (*Il-6*) and Brown-Forsythe and Welch ANOVA (*Il-18, Ifn-γ, Tnf-α, Il-10, p65, Ctgf, Fibronectin, Tgf-β, Mmp2, Mmp9 and Gsdmd*). *n*=7-13 per group. *P<0.05, **P<0.01, ***P<0.001, ^#^P<0.05, ^##^P<0.01, ^###^P<0.001 and ^####^P<0.0001 as indicated or vs I/R group. Results are expressed as mean ± SEM.

At 28 days post-AMI, *Il-6, Il-18* and *Ifn-γ* were elevated by 3.6-, 3.6- and 2.5-fold respectively compared to their sham control groups. Importantly, at this timepoint, treatment with DSF reduced the gene expression of all inflammatory markers assessed (Figure 6B and SFigure 2C). Interestingly, the expression of the anti-inflammatory cytokine, *Il-10* also increased by 3.9-fold (Figure 6B) in the vehicle-treated I/R group compared with the sham group and was reduced by DSF (50mg/kg) treatment.

At 7 days post-AMI, the expression levels of fibrosis genes such as *Ctgf,* fibronectin and *Tgf-*β were significantly increased (Figure 6C) compared to the sham group, while *Mmp2* and *Mmp9* expression remained unaltered. Importantly, there was a significant reduction in *Ctgf*, *fibronectin* and *Tgf-β* gene expression (Figure 6C) with DSF (50mg/kg) treatment. At the 28-day post-AMI timepoint, the expression of *Ctgf, Fibronectin*, *Tgf-*β and *Mmp2* was significantly elevated (Figure 6D) compared to the sham group, whilst MMP9 showed no change. A highly significant reduction in *Ctgf*, *Fibronectin*, *Tgf-*β, and *Mmp2* was observed after DSF treatment (50mg/kg) (Figure 6D) compared to the vehicle-treated I/R group and importantly, these reductions were evident at both DSF concentrations.

At 7 days post I/R injury, inflammasome and cell death-related gene expression levels of *p65* (NF-κβ subunit) and *Nlrp3* were not significantly elevated (Figure 6E). However, the gene expression levels of *Caspase-3* and *Gsdmd* showed a trend toward increased expression at this time point. At 28 days post-AMI, *Nlrp3, Gsdmd* and *Caspase-3* levels were significantly elevated by 1.6-; 1.6-, and 1.8-fold respectively compared to the sham group (Figure 6F). Importantly, at 7 days post-AMI, *Caspase-3*, *p65*, *Gsdmd* and *Nlrp3* gene expression levels were reduced by DSF (50mg/kg) treatment (Figure 6E). At 28 days post-AMI, DSF (both 25mg/kg and 50mg/kg) treatment significantly attenuated *Caspase-3, p65, Gsdmd* and *Nlrp3* gene expression as compared to vehicle-treated I/R groups (Figure 6F).

The ROS producing NADPH oxidase gene, *Nox4*, was significantly elevated at both the 7-day and 28-day timepoints (3.8- and 8-fold respectively, SFigure 2B and SFigure 2D) post-I/R injury compared to its corresponding sham group, whilst DSF treatment (50mg/kg) led to a significant reduction in *Nox4* gene expression compared to the vehicle-treated I/R groups (SFigure 2B and SFigure 2D).

### 3.7 Effect of disulfiram on cardiac proteins related to cell death and the NLRP3-Gsdmd axis

The changes observed in the gene expression of *Nlrp3, Gsdmd* and *Caspase-3* were reflected in the changes in their protein levels in heart tissue at 28 days post-I/R injury. The levels of GSDMD-full length (FL), GSDMD-C terminal (CT), Caspase-1 and Caspase-3 were significantly elevated post-AMI compared to sham controls (Figure 7). DSF treatment led to significant reductions in GSDMD-CT and Caspase-1 and -3, at both 25- and 50mg/kg DSF, whilst GSDMD-FL showed a significant reduction at 25mg/kg (Figure 7A and 7C). Processed Caspase-1 (p10/12) and Caspase-3 (p17) were not detected. There was a tendency towards an increase in the NLRP3 protein which was reduced with DSF treatment, although the results were not significant (Figure 7A and 7B). With respect to GSDMD cleavage, the band observed at ∼30kDa is most likely the pore-forming GSDMD-NT fragment which was not highly expressed. In addition, the pattern of expression of the GSDMD-NT fragment mirrored the expression of the GSDMD-CT fragment. Moreover, there is the potential for Caspase-3-mediated cleavage of GSDMD as indicated by the appearance of a p43 fragment, although this cannot be validated without the appropriate controls^21^.

**Figure 7:**
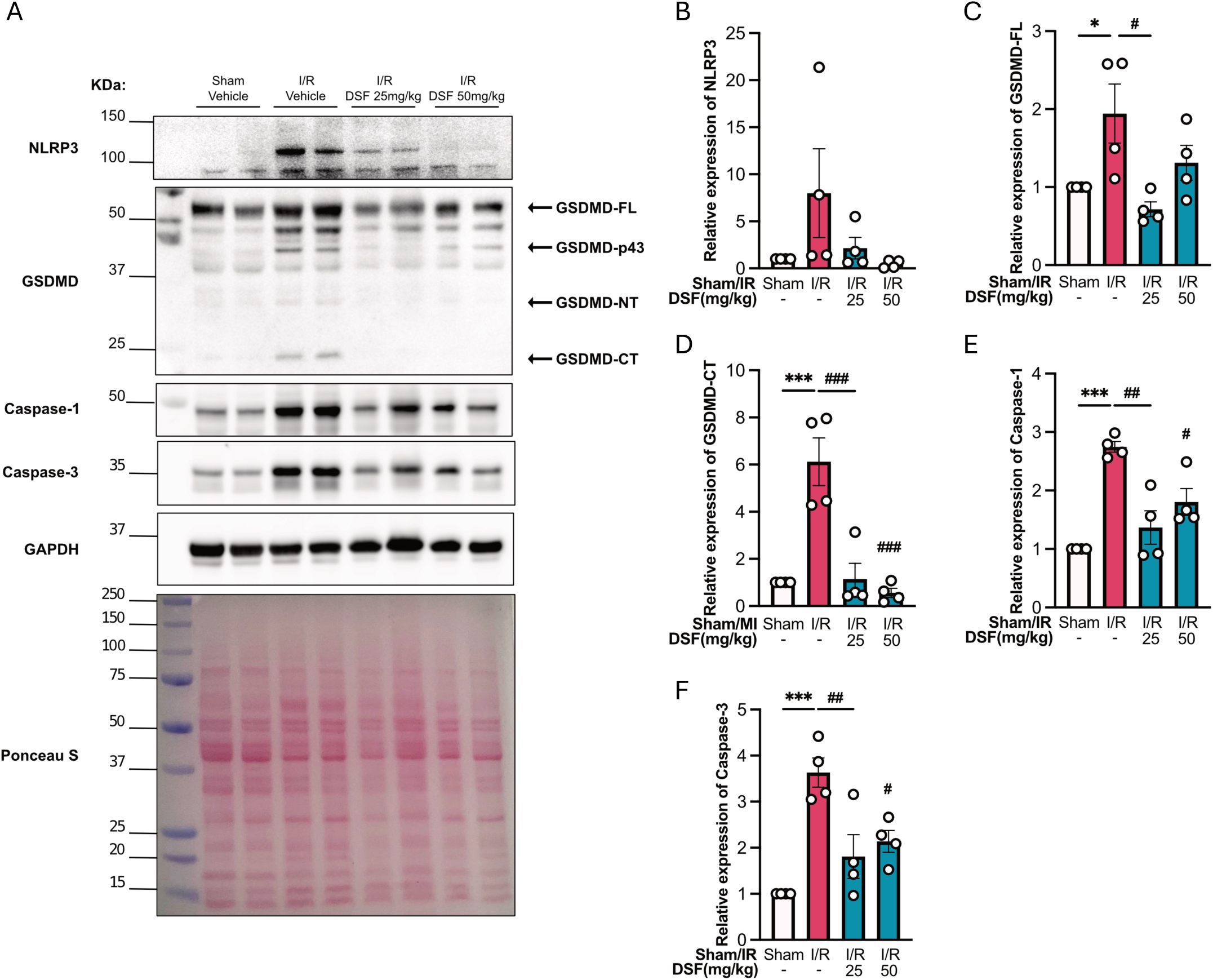
(A) Immunoblot and quantification of (B) NLRP3, (C) GSDMD-FL, (D)GSDMD-CT, (E) Caspase-1 and (F) Caspase-3 in heart 28 days post-I/R injury treated with vehicle or DSF. Total proteins are normalised against GAPDH and analysed using Ordinary one-way ANOVA. *n* represents the number of biological replicates, which are graphed as individual points. *n*=4. *P<0.05, ***P<0.001, ^#^P<0.05, ^##^P<0.01 and ^###^P<0.001 as indicated or vs I/R group. Results are expressed as mean ± SEM.

## 4. Discussion

This study evaluates the therapeutic potential of the FDA-approved drug DSF, recently identified to have anti-inflammatory properties, to determine whether inhibition of GSDMD pore formation alleviates immune-mediated inflammation and cardiac injury post-AMI. We show that DSF reduces IL-1β secretion in both human and mouse macrophage *in vitro* and decreases immune cell presence in murine cardiac tissue *in vivo*. Mechanistically, DSF treatment reduced pro-inflammatory gene and protein expression. Consistent with reduced inflammatory activation, DSF treatment also decreased IL-6 secretion from BMDMs, indicating a coordinated suppression of pro-inflammatory cytokine production. Our temporal *in vivo* analysis post-AMI revealed significant structural improvements by DSF, with early functional improvements in ejection fraction. Thus overall, our data support a strong anti-inflammatory role for DSF that reduces cardiac fibrosis and improves cardiac function.

The GSDMD pore-forming protein has attracted attention for its pivotal role in mediating inflammation and pyroptosis. Accordingly, GSDMD pore formation is implicated in disease, including ischemic stroke, sepsis, cancer and viral infections^13, 19^. Secreted IL-1β is the major pro-inflammatory cytokine that drives IL-6 production and its release from inflammatory cells is commonly mediated via GSDMD, however it can also occur via GSDMD-independent mechanisms^19^. Therapeutic inhibition of IL-1β gained support from the CANTOS trial, which reduced adverse CV events after AMI,^22^ although increased fatal infections highlighted the need to fine-tune the reduction in IL-1β while preserving host defences against infections. Thus, targeting GSDMD post-AMI to limit IL-1β-mediated cytotoxicity offers an alternate strategy, yet enables the release of Il-1β via GSDMD-independent mechanisms. However, whether GSDMD-independent IL-1β secretion from resting or non-pyroptotic cells such as neutrophils or cells that do not express GSDMD, is sufficient to limit infections remains unclear^23^.

DSF was identified as an effective inhibitor of GSDMD pore formation via high-throughput biochemical screening conducted by Hu et al^13^. In our study, using both ELISA and immunoblot analysis, we showed that DSF significantly attenuated IL-1β secretion in a dose-dependent manner (0.1-50 µM) from both human and murine macrophages and provided protection against lytic cell death, also known as pyroptosis. Additional mechanistic insight was obtained by immunoblotting, where DSF reduced both the production of processed GSDMD-NT protein and that of the pro- and secreted forms of IL-1β, with no effect on Caspase-1. This suggests that in addition to the known inhibitory effect of DSF on GSDMD via direct binding, DSF also inhibits the processing of GSDMD into its active NT fragment in a dose-dependent manner, although this appeared to be independent of changes in Caspase-1.

This study shows that DSF reduces IL-6 transcription and secretion, suggestive of a feed-forward mechanism to regulate inflammatory responses. This was evidenced by the decrease in the gene expression of several pro-inflammatory mediators such as *Tnf-α* and *Mcp-1,* in agreement with previous findings^24^.

In our current study, we evaluated the effects of DSF in an I/R injury animal model which more closely mimics the clinical scenario where heart attack patients receive revascularisation via percutaneous coronary intervention (PCI) or, in severe cases, coronary artery bypass surgery (CABG) to restore blood flow^25^. The re-introduction of blood into an ischemic zone generates ROS, an influx of Ca2+, and an alkaline pH of the blood (alkalosis), which are all known drivers of inflammation^5, 26, 27^. Our rationale for testing DSF was two-fold. DSF was identified as a strong inhibitor of inflammatory GSDMD pore formation, yet its potential to limit inflammasome-driven cytokine release and downstream cardiac injury has not been fully explored pre-clinically in a context-relevant animal model. Furthermore, as an FDA-approved compound with an established safety profile, DSF offers an opportunity for drug repurposing.

To reflect the clinical context where patients receive treatment following a PCI or CABG, DSF was administered immediately upon reperfusion, and continued daily until termination. This study then assessed changes in immune cell populations within the heart, spleen, bone marrow and blood to determine whether DSF treatment modulates immune cell populations and mobilisation in response to cardiac I/R injury. Our study revealed an increase in leukocytes in the heart post-AMI, as evidenced by the rise in the pan-leukocyte marker CD45 on flow analysis and CD68 via IHC, indicative of increased macrophages in the myocardium. This was evident at 7 and 28 days post-AMI, and aligns with previous findings of an early elevated and ongoing presence of leukocytes post-I/R injury^28^. There was also an increase in *Mcp-1* gene expression 28 days post-AMI, which is known to promote the recruitment of monocytes/macrophages into the infarcted heart, inducing cytokine production and triggering the inflammatory response^29^. DSF treatment caused a reduction of monocyte infiltration as well as CD68 expression in the heart, particularly within the infarct region at 7 days post-AMI, and attenuated MCP-1 gene expression at 28 days post-AMI. Taken together, our data strongly suggests that DSF mediates the reduction in monocytes and macrophages within the infarcted heart, which in turn attenuates the ongoing inflammation by decreasing the secretion of pro-inflammatory cytokines.

From a mechanistic perspective, during the cardiac remodelling phase of an AMI, a phenotypic switch occurs in which monocytes switch to an anti-inflammatory Ly6C-low subset to reduce inflammation and promote healing^30^. In our study, DSF treatment caused a reduction in both Ly6C-high and Ly6C-low subsets in the spleen, whilst a phenotypic switch toward Ly6C-low monocytes occurred in the bone marrow. Although there was no overall difference in total number of monocytes in the bone marrow, DSF treatment not only reduced the number of pro-inflammatory Ly6C-high monocytes, but led to the expansion of the anti-inflammatory Ly6C-low monocyte population. Importantly, it is known that increased infiltration and persistence of Ly6C-high monocytes and depletion of Ly6C-low monocytes correlate with reduced cardiac function^30, 31^. Hence, this phenotypic switch towards the Ly6C-low monocytes in the bone marrow and an overall reduction of monocytes in the spleen, a known reservoir for cardiac leukocytes post-AMI^32^, likely contributed to the reduction in leukocyte-mediated cardiac injury by DSF seen in this study. Collectively, DSF appears to attenuate the self-perpetuating cycle of leukocyte-mediated inflammation to ultimately alleviate cardiac injury post-AMI.

GSDMD-mediated pyroptosis has been demonstrated to be the dominant form of cell death responsible for left ventricular remodelling and cardiac dysfunction post-AMI^33^. Pyroptosis is a highly inflammatory-mediated lytic form of cell death, resulting from the priming and activation of the NLRP3-inflammasome, increasing plasma membrane permeability and facilitating the release of inflammatory cytokines. In the current study, NLRP3, GSDMD and Caspase-1, the principle drivers of pyroptosis, were upregulated in the heart 28 days post-AMI, suggesting ongoing activation of the pyroptotic pathway in this model which is consistent with the literature^9, 33^. Importantly, DSF treatment significantly reduced their expression levels at 7 and 28 days post-AMI in a dose-dependent manner. Similarly, the apoptosis mediator, *Caspase-3,* was upregulated on days 7 and 28 days post-AMI, and DSF treatment abrogated these increases. This indicates that DSF induces early changes in gene and protein expression of key pyroptosis and apoptosis-related mediators to promote a decrease in cell death, thereby positively contributing to reduced cardiac injury. Furthermore, as cardiomyocytes are terminally differentiated cells with minimal regenerative capacity, prevention of cell death is considered essential in the treatment of CVD^34^.

ROS are potential activators of the NLRP3 inflammasome and contribute to elevated inflammation and cell death^35^. Nox4 is prominent in post-AMI injury as it induces cardiac hypertrophy, fibrosis and apoptosis in response to MI^36^. Moreover, inhibition of Nox2 and Nox4 have been shown to improve cardiac function post-AMI^37^. In our study, DSF reduced elevated cardiac *Nox4* gene expression at both 7 and 28 days post-AMI. By inference, the reduction in *Nox4* following DSF treatment may contribute to a decrease in cardiac hypertrophy^38^, as observed by the lower heart weight and LV weight-to-tibia length ratio at 7 days post-AMI in this study. Reducing cardiac hypertrophy helps lessen the stress on the heart and prevents further cardiac injury, thereby aiding in the preservation of cardiac function by limiting infarct expansion and myocardial rupture^38^.

The primary aim of therapies after a heart attack is to prevent HF, a common and often fatal complication affecting roughly half of all patients^4, 39, 40^, and driven by extensive fibrosis. In the current study, well-known fibrotic markers as *Ctgf*, *Fibronectin* and *Tgf-β* were elevated at 7 and 28 days post-AMI. Treatment with DSF reduced these markers in the heart at both 7 and 28 days post AMI, which correlated with a decrease at the transcriptional level. Our data are therefore highly suggestive of an anti-fibrotic effect by DSF within the heart in the aftermath of an AMI. Increased fibrosis post-AMI reduces LV contractility which in turn increases the pressure to compensate for the loss of viable tissue, causing increased wall stress and pressure overload^41^. Over time, pressure overload can lead to HF as the heart struggles to cope with the elevated pressure^41^. Hence, reducing cardiac fibrosis is essential to minimise the area of damaged cardiac tissue to reduce the infarct size and preserve LV contractility. Ultimately, this reduces the risk of HF. Inhibition of CTGF is known to attenuate LV systolic dysfunction and lessen inflammatory and pro-fibrotic gene expression [52]. On the other hand, TGF-β has both reparative and inflammatory roles. In the early phase, TGF-β can reduce myocardial injury, but prolonged expression leads to the active conversion of fibroblasts into myofibroblasts and causes scar formation [53]. In addition, DSF reduced *Mmp2* gene expression, a known driver of fibrosis, at 28 days post-AMI, potentially attenuating fibrotic and inflammatory remodelling through decreased MMP2 activity. This further aligns with our finding that cardiac fibrosis was reduced at both 7 and 28 days post-AMI after drug treatment.

Cardiac fibrosis post-AMI significantly alters systolic function as the heart undergoes dilative remodelling and experiences inadequate filling and contraction. In our study, DSF improved systolic function as seen by improvements in diastolic and systolic volume and area, and an improvement in ejection fraction at 7 days post-AMI, albeit that LV shortening remained unaltered. Ejection fraction correlates with myocardial viability^42^. Furthermore, an improvement in ejection fraction attenuates long-term pressure overload by facilitating the ejection of a larger volume of blood with each contraction, thereby reducing the pressure in the LV. This alleviates the strain caused by sustained pressure overload, which lessens cardiac remodelling and HF^43^. Hence, it can be postulated that DSF improved early cardiac function via reductions in pressure overload in the LV, as seen by improvements in EF at 7-day post-AMI. From a mechanistic perspective, the improvement in early cardiac function correlated with reductions in leukocyte infiltration into the myocardium, the promotion of an anti-inflammatory phenotype and a reduction in cardiac fibrosis, suggesting that underlying structural changes may have driven functional improvements. However, despite the positive effects of DSF on cardiac structure at 28-days post AMI, improvements in cardiac function, particularly ejection fraction, were not evident at this time-point. First, it is possible that a dose of 50 mg/kg was insufficient to achieve long-term improvements in cardiac function. Second, a DSF-mediated improvement in ejection fraction might be masked by the action of pressure overload in control mice post-AMI, resulting in similar ejection fraction levels at the later time-point. This is postulated because of the DSF-mediated reduction in cardiac fibrosis at 28 days post-AMI, independent of improvements in cardiac function. The reduction in cardiac fibrosis would ensure that contractility is preserved, preventing the heart from overworking to compensate for the loss of viable cardiac tissue in maintaining the same level of ejection fraction. This reduces the risk of HF and aligns with the literature where patients that experience HF with preserved ejection fraction, show high left ventricular pressures and impaired ventricular relaxation, despite normal or near normal ejection fraction^44^. Future studies should assess measures of pressure overload by pressure-volume catheterisation to determine if DSF treatment improves this parameter, which is an important predictor of HF.

## 5. Conclusion

In summary, we show the anti-inflammatory effect of DSF in modulating IL-1β secretion from inflamed cells as an effective means to halt the inflammatory cascade that contributes to adverse cardiac remodelling post-AMI. DSF treatment resulted in reduced pro-inflammatory cytokine expression, which further attenuated the production and infiltration of immune cells into the heart, thereby decreasing the death of cardiac tissue and alleviating cardiac fibrosis. A reduction in cardiac fibrosis ultimately aided in preserving cardiac function, as evident 7 days post-AMI. In addition, dosage-dependent changes in gene expression were observed after DSF treatment, suggesting that careful dosage considerations may be needed before moving DSF forward into the clinic for the treatment of post-AMI HF, albeit that this was only assessed at the transcriptional level. For example, it is interesting that at 7 days post-AMI, the reduction in pro-inflammatory and pro-fibrotic gene expression was only evident at the higher dosage of DSF (50 mg/kg), while both low (25 mg/kg) and high (50 mg/kg) doses of DSF similarly attenuated pro-fibrotic and inflammatory gene expression at 28 days post-AMI. This may be attributed to the duration of treatment where a shorter treatment time (7-days post I/R-injury) required the higher dosage given the intense inflammatory response in the immediate aftermath of an AMI. This may be the reason why previous studies that used DSF to attenuate inflammation mainly used the higher dose (50mg/kg) of DSF^13, 14, 45^ for shorter duration^14^. With respect to its safety profile, DSF is known to modify additional cellular proteins and this could result in off-target effects^13^. However, the well-established safety profile of DSF in humans, demonstrated over six decades of clinical use for alcoholism, suggests that modifications to other cellular targets do not result in significant clinical toxicity. Moreover, the transcriptional impact of DSF treatment, as well as its ability to target GSDMD, may confer added benefit compared to GSDMD-targeting alone. Therefore, these findings pave the way for repurposing DSF as a potent cardioprotective agent, and for the development of more specific GSDMD inhibitors for post-AMI therapy. Overall, our findings suggest that targeting post-AMI inflammation with DSF not only attenuates cardiac injury but also facilitates the recovery of cardiac structure and function.

## 6. Translational Perspective

In recent years, mortality trends after AMI have changed little, highlighting the urgent need for better treatment options. Whilst several anti-inflammatory therapies were effective in reducing cardiovascular risk after MI, they were often associated with a higher risk of fatal infections. Hence, fine-tuning of inflammation is necessary. We preclinically demonstrated that repurposing the FDA-approved drug Disulfiram to inhibit GSDMD can attenuate inflammation after I/R injury following MI. This resulted in the significant reduction of cardiac inflammation and fibrosis after I/R injury after DSF treatment which preserved cardiac function. These findings support Disulfiram and GSDMD inhibition as a translational pathway from inflammation to cardioprotection in patients with MI.

## 8. Acknowledgement

The authors thanks the Baker Institute Flow Cytometry, the Alfred Research Alliance Animal Ethics Committee and the Monash Histology Platform.

## 9. Author contribution statement

J.S.C., A.S. and J.B.D.H. designed research studies and wrote the manuscript. J.S.C., A.S., D.D., H.K. and A.B participated in the animal experiment and checked the animal data. J.S.C., D.D., M.P., H.K. and M.K.S.L. conducted, acquired, and interpreted data. M.P and P.Y participated in the animal experiment. A.J.M. and J.E.V. contributed to the study design and supervised the project.

## 11. Data availability statement

The datasets used and/or analysed during the current study are available from the corresponding author on reasonable request.

