## Supplemental Figure for "Inhibition of Gasdermin D by Disulfiram Attenuates Cardiac Inflammation and Fibrosis following Ischaemia Reperfusion Injury"

^7^Co-senior

**Short title: Targeting Gasdermin D improves Cardiac Remodelling after I/R injury**

**Addresses for correspondence:**

**Prof. Judy de Haan**

Lab Head (Cardiovascular Inflammation and Redox Biology Laboratory)

Heart Failure Program

Baker Heart and Diabetes Institute

75 Commercial Road

Melbourne, Victoria 3004, Australia

***Supplementary figures***

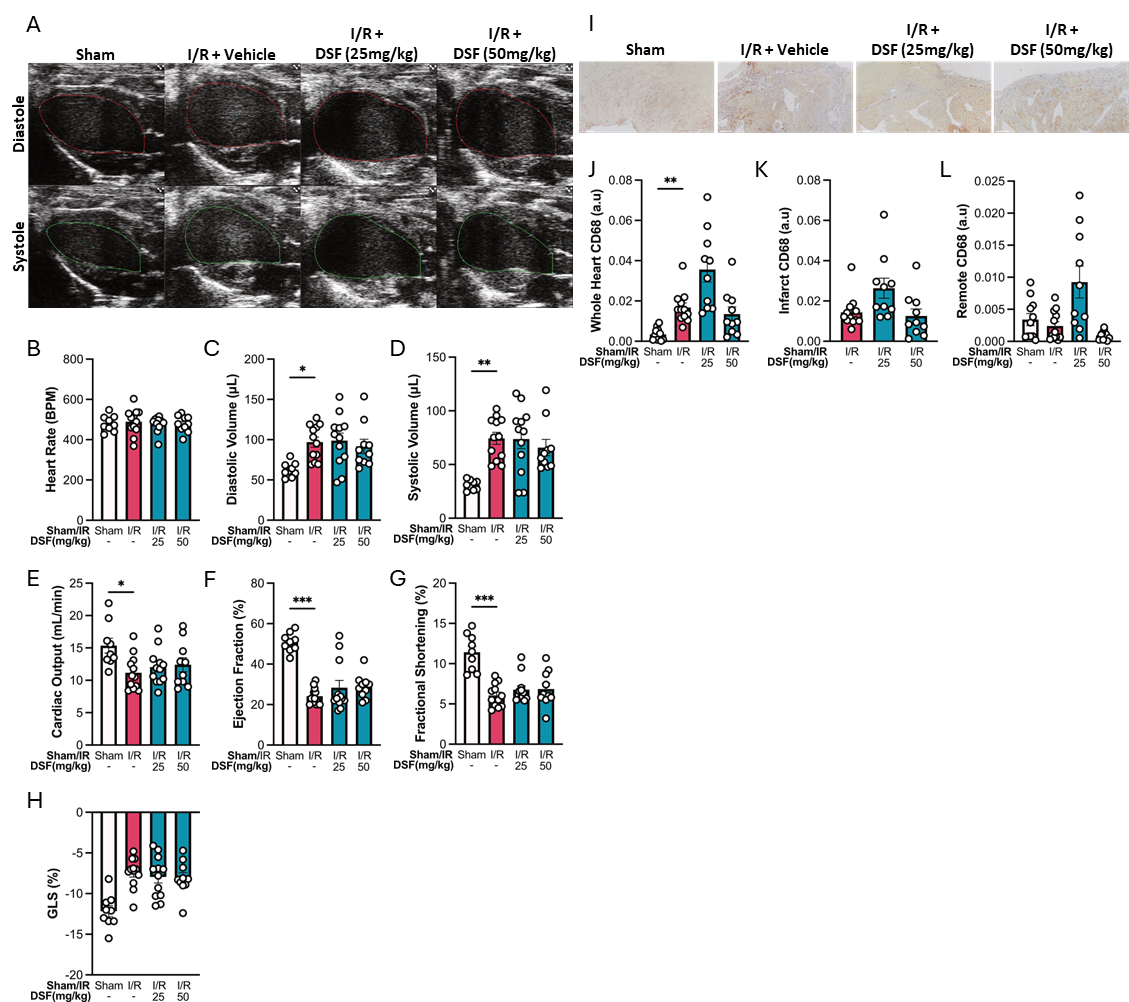

**Supplementary Figure 1:**

(A) Representative echocardiographic images at end-phase diastole and end-phase systole. The red lines show the tracing of the endocardium at maximum relaxation (diastole) and the green lines reflect maximum contraction at systole. Analysis of (B) Heart rate, (C) Diastolic volume, (D) Systolic volume, (E) Cardiac output, (F) Ejection fraction, (G) Longitudinal fractional shortening (H) Global longitudinal strain in sham and I/R mice treated with vehicle or DSF 28 days post-I/R injury. (I) Representative images of CD68 stained hearts and quantification in the (J) whole heart, (K) infarct region and (L) remote region 28 days post-I/R injury treated with vehicle or DSF. Scale bars represent 2mm. *n* represents the number of biological replicates, which are graphed as individual points. Statistics used: Ordinary one-way ANOVA (C, E) and Kruskal-Wallis (D, F, G, J). n= 9-12 per group. *P<0.05, **P<0.01, ***P<0.001. Results are expressed as mean ± SEM.

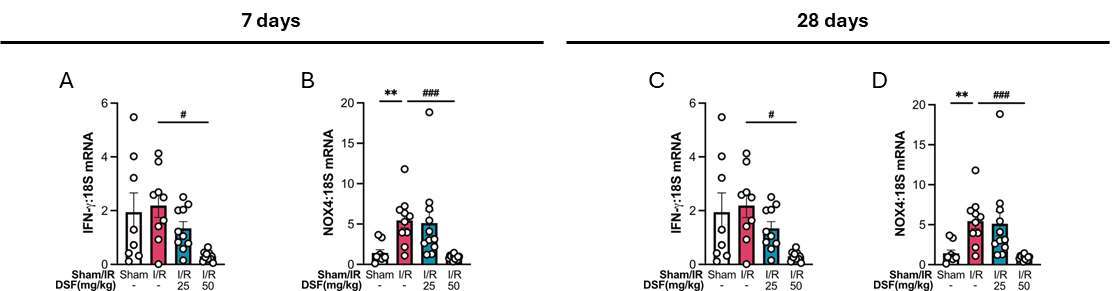

**Supplementary Figure 2:**

Gene expression of (A) *Ifn-γ* and (B) *Nox-4* at 7 days, and (C) *Ifn-γ* and (D) *Nox-4* at 28 days post-I/R injury in the heart with vehicle or DSF treatment measured by qRT-PCR. Gene expression is shown relative to 18S mRNA levels. n represents the number of biological replicates, which are graphed as individual points. Statistics used: Brown-Forsythe and Welch ANOVA (A, D), Kruskal-Wallis (B) and Ordinary one-way ANOVA (C). *n*=5-12 per group. **P<0.01, ^#^P<0.05 and ^###^P<0.001. Results are expressed as mean ± SEM.

**Supplementary Table 1:**

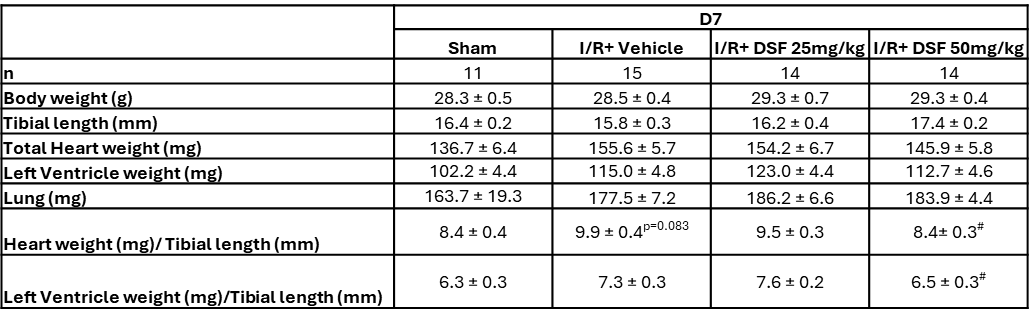

Morphometric analysis at the conclusion of 7 days post-I/R injury. Statistics used: Ordinary one-way ANOVA (Heart weight/tibia length and Kruskal-Wallis (LV weight/tibia length). n=11-15. ^#^P<0.05 vs I/R + vehicle. Results are expressed as mean ± SEM.

**Supplementary Table 2:**

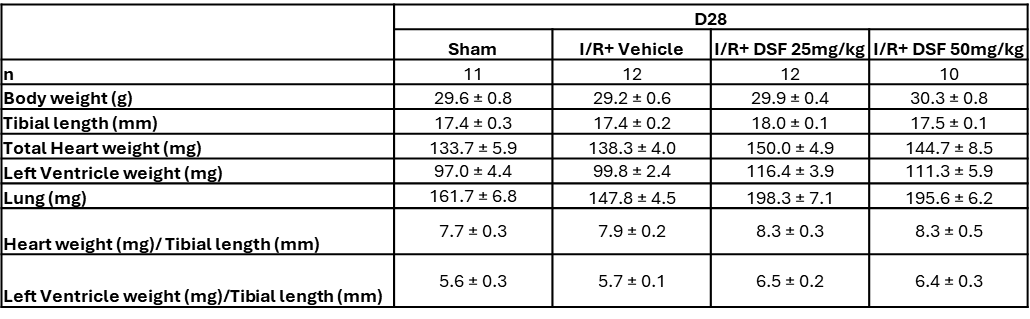

Morphometric analysis at the conclusion of 28 days post-I/R injury. n=10-12.

**Supplementary Table 3:** The complete list of antibodies/dyes used in the study and their working concentrations.

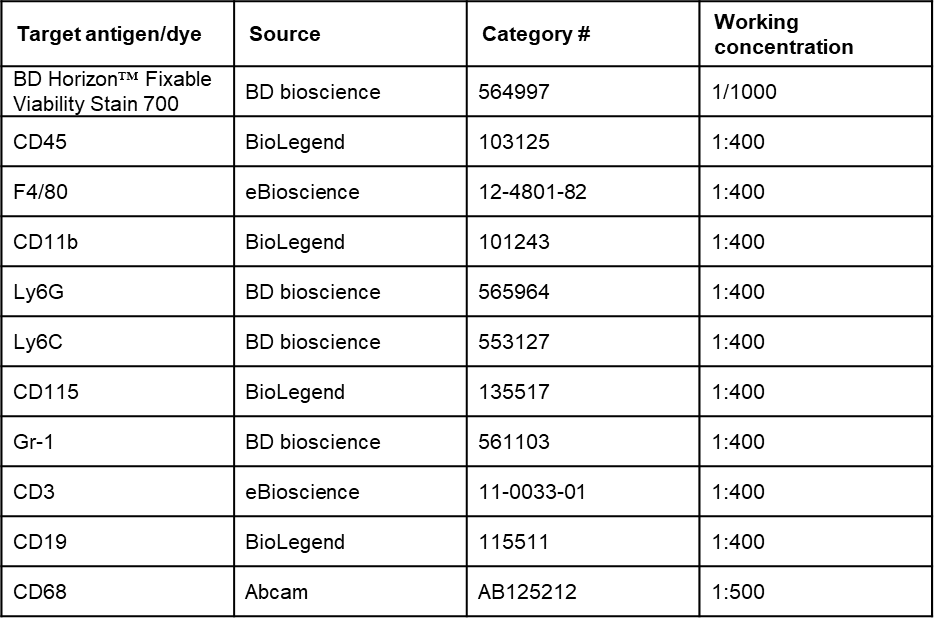

**Supplementary Table 4:** The complete list of primer sequences used in the study for gene expression analysis by qRT-PCR.

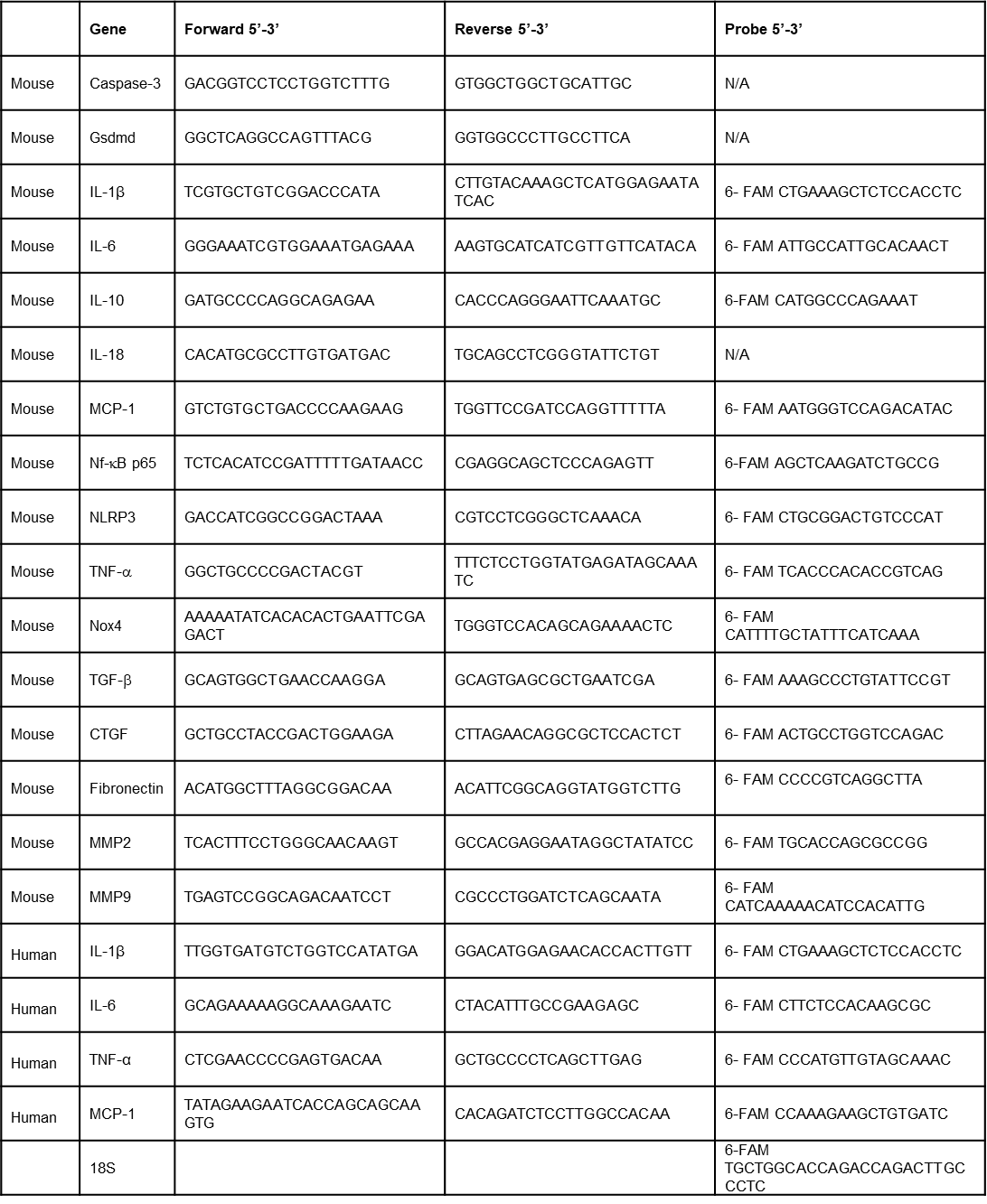
